# The homolog of human ecto-nucleoside triphosphate diphosphohydrolase MIG-23 is involved in sperm migration by modulating extracellular ATP levels in *Ascaris suum*

**DOI:** 10.1101/2021.12.14.472724

**Authors:** Qiushi Wang, Ruijun He, Lianwan Chen, Qi Zhang, Jin Shan, Yanmei Zhao, Xia Wang

**Affiliations:** Key Laboratory of RNA Biology, CAS Center for Excellence in Biomacromolecules, Institute of Biophysics, Chinese Academy of Sciences, Beijing 100101, China; University of Chinese Academy of Sciences, Beijing 100049, China; National Laboratory of Biomacromolecules, CAS Center for Excellence in Biomacromolecules, Institute of Biophysics, Chinese Academy of Sciences, Beijing 100101, China

**Keywords:** sperm migration, adenosine-5’-triphosphate (ATP), MIG-23, *Ascaris suum*

## Abstract

In nematodes, spermiogenesis, which is also called sperm activation, is a process in which nonmotile spermatids are transformed into crawling spermatozoa, which is accompanied by a series of morphological, physiological and biochemical changes. Sperm motility acquisition during this process is essential for successful fertilization. However, the mechanisms of sperm motility regulation in nematodes remain to be clarified. Herein, we found that extracellular adenosine-5’-triphosphate (ATP) level mediation by MIG-23, which is a homolog of human ecto-nucleoside triphosphate diphosphohydrolase (E-NTPDase), was required for major sperm protein (MSP) filament dynamics and sperm motility in the nematode *Ascaris suum*. MIG-23 was localized on the sperm plasma membrane. During sperm activation, mitochondrial activity was increased dramatically, and a large amount of ATP was produced and stored in refringent granules (RGs). In addition, a portion of the produced ATP was released to the extracellular space through ATP channels, which were composed of innexins and localized on the sperm plasma membrane. Spermatozoa, instead of spermatids, hydrolyzed exogenous ATP and processed ecto-ATPase activity. MIG-23 contributed to the ecto-ATPase activity of spermatozoa. Once MIG-23 activity was interrupted, spermatozoa also decreased their ATP hydrolysis activity. Blocking MIG-23 activity resulted in an increase in the depolymerization rate of MSP filaments in pseudopodia, which eventually affected nematode sperm migration. Overall, our data imply that MIG-23, which contributes to the ecto-ATPase activity of spermatozoa, regulates sperm migration by modulating extracellular ATP levels.

**Highlights:**

1. ATP is released to extracellular space through innexin channels which are identified in worm sperm.
2. Worm spematozoa show ecto-ATPase activity.
3. MIG-23 contributes to the ecto-ATPase activity of spermatozoa and regulates extracellular ATP level.
4. MIG-23 is required for MSP-based filament dynamics and sperm migration.

## Introduction

Unlike mammalian sperm, which are flagellated via the tubulin cytoskeleton, nematode sperm crawl via the major sperm protein (MSP)-based cytoskeleton. Nematode sperm migration requires both protrusion of the leading edge and retraction of the cell body, coupled with adhesion to the substrate (Prass et al., 2006). Previous studies have demonstrated that the assembly and disassembly of MSP filaments regulate sperm motility directly. The dynamics of MSP filaments are modulated by factors such as membrane tension, MSP filament-associated proteins and phosphylation level. Sperm membrane tension regulates cell migration by controlling MSP-based cytoskeletal dynamics in *Caenorhabditis elegans* sperm (Batchelder et al., 2011). MSP-associated protein phosphorylation and dephosphorylation regulate nematode sperm motility (Yi et al., 2009). Sperm-specific PP1 phosphatases (GSP-3/4), which are spatial regulators of MSP disassembly, modulate sperm motility in *Caenorhabditis elegans* (Wu et al., 2012). In addition, intracellular Ca^2+^, cholesterol and the biosynthesis of glycosphingolipids were found to be required for nematode sperm migration (Dou et al., 2012a; Shang et al., 2013). Here, we found that regulation of the extracellular ATP level by MIG-23, which is a homolog of ectonucleoside triphosphate diphosphohydrolase (E-NTPDase), is required for sperm migration in *Ascaris suum*.

ATP, which is an energy molecule or signaling molecule, is required for various physiological functions, including cellular metabolism, cell adhesion, activation, proliferation, differentiation and migration (Fujii et al., 2012). ATP produced by mitochondria is released from cells into the extracellular space via stimulated exocytosis and ATP channels (Lazarowski, 2012; Praetorius and Leipziger, 2009). The released ATP plays a role in cell functions. Human keratinocytes release ATP, and the released ATP acts as feedback to activate its receptors to further elevate the intracellular calcium concentration (Ho et al., 2013). Released ATP also regulates various key physiological processes. For instance, it modulates central synaptic transmission in neurons and microglial migration (Dou et al., 2012b; Zhang et al., 2003). In ATP release-defective transgenic mice, synaptic cross-talk is affected and results in heterosynaptic depression (Pascual et al., 2005). Both ATP release and ATP hydrolysis are involved in neutrophil chemotaxis (Chen et al., 2006; Corriden et al., 2008a). Extracellular ATP in the female reproductive tract facilitates the bovine spermatozoa acrosome reaction, and this process favors sperm fusion with oocytes (Luria et al., 2002). Intracellular ATP is released into the extracellular space via ATP channels and exocytosis (Dou et al., 2012b; Zhang et al., 2007). ATP that is stored in lysosomes is released through exocytosis in astrocytes (Zhang et al., 2007). Connexin and pannexin are hemichannels that mediate ATP release (Romanov et al., 2007). On the astrocyte cell surface, ATP release channels are colocalized with ecto-ATPase (Joseph et al., 2003). Furthermore, the E-NTPDase family is responsible for hydrolyzing extracellular ATP to ADP and AMP. E-NTPDases control the concentrations of extracellular nucleotides. Eight members of the E-NTPDase family have been identified in mammals. NTPDase1, NTPDase2, NTPDase3 and NTPDase8 are localized on the plasma membrane. These membrane-bound NTPDases are glycoproteins and are present in various tissues, such as the brain, lung, heart, and kidney. NTPDase1 was detected in the testis, and NTPDase6 was putatively localized in acrosome vesicles of round spermatids in mice (Martín-Satué et al., 2009). NTPDase1, NTPDase2 and NTPDase8 were localized in the rat uterus (Milošević et al., 2012). NTPDases are involved in multiple and diverse physiological processes, such as pathogen-host interactions, lipid glycosylation, eye development, and taste bud function (Masse et al., 2007; Schachter et al., 2015; Vandenbeuch et al., 2013). Rat NTPDase1 inhibits platelet aggregation and favors blood flow (Zebisch et al., 2012). Deficiency of E-NTPDase1 in mice was found to affect male fertility by reducing the concentration of spermatozoa in the semen (Kauffenstein et al., 2014). Ecto-ATPases, which determine extracellular ATP homeostasis, are present in cells of various types and modulate important physiological functions. MIG-23 in *Caenorhabditis elegans*, which is an NDPase that interacts with MIG-17, an ADAM (a disintegrin and metalloprotease), regulates distal tip cell (DTC) migration (Nishiwaki et al., 2004). Here, we found that in the nematode *Ascaris suum*, MIG-23, which is a homolog of human NTPDase, is involved in sperm migration by modulating extracellular ATP levels.

## Results

### Plentiful ATP is produced, and a portion of the produced ATP is stored in refringent granules during nematode sperm maturation

Sperm maturation in *Ascaris suum* is associated with the formation of a motile pseudopodium, fusion of membranous organelles (MOs) with the plasma membrane and coalescence of refringent granules (RGs) (Abbas and Cain, 1981). In addition to these changes in morphology and physiology, we found that the mitochondrial membrane potential (MMP) increased significantly during sperm maturation. The MMP reflects the mitochondrial activity (Lachance et al., 2013). It was monitored with a fluorescent indicator, namely, JC-1, which was used to measure the MMP of spermatids (nonmature) and spermatozoa (mature). As shown in Fig EV1A, compared to that in spermatids, the MMP level in spermatozoa was much higher. The MMP of spermatozoa increased nearly 3-fold compared with that of spermatids (Fig 1A). This indicates that spermatozoa generate a much greater supply of ATP to meet needs such as protein phosphorylation, plasma membrane extension, vesicle fusion, and cytoskeleton assembly. To measure cytosolic ATP levels during sperm maturation, ATP concentrations were measured using a luciferin-luciferase assay. The results showed that cytosolic ATP levels rose at the beginning of sperm maturation (~5 min) and subsequently declined (~15-30 min) (Fig 1B). However, at the same time point, spermatozoa maintained a higher MMP (~15-30 min) (Fig EV1B). This raises an interesting issue as to why mitochondrial activity is higher but the cytosolic ATP level is lower. Three possible pathways may result in a lower ATP level in the cytosol. First, physiological activities, such as cytoskeleton assembly, protein phosphorylation and sperm motility, consume ATP during sperm maturation (Miao et al., 2003). Second, some ATP may be stored in vesicles. Third, some ATP may be released into the extracellular space. We tested whether nematode sperm are able to store ATP. Quinacrine is an ATP storage marker and indicates ATP localization in cells (Haanes et al., 2014). We found that quinacrine accumulated in dispersed vesicles in spermatids. In spermatozoa, quinacrine accumulated in a large organelle (Fig 1C). The quinacrine intensity of spermatozoa was stronger than that of spermatids. Flow cytometry analyses also demonstrated that the fluorescence intensity of quinacrine in spermatozoa was more than 2-fold higher than that in spermatids (Fig 1D). The dynamic process of ATP storage was consistent with RG fusion. The quinacrine intensity of sperm increased significantly during sperm maturation. The process was observed and images were obtained with a confocal microscope (Fig 1E, Movie EV1). Next, we tested the possibility of ATP release. The released ATP (in medium) of spermatids and spermatozoa that were activated at the 5th minute was measured. Compared to the control, the ATP concentration in the medium increased significantly in spermatids and spermatozoa (Fig 2D). This indicates that both spermatids and spermatozoa are able to release ATP from the cytosol into the extracellular space. In addition, we tested whether ATP that is produced by mitochondria is required for sperm maturation. Inhibitors of mitochondrial function, such as CCCP (an uncoupler of oxidative phosphorylation), antimycin (an inhibitor of cellular respiration, specifically oxidative phosphorylation), and oligomycin (an inhibitor of ATP synthase), were used to treat sperm. All these reagents inhibited ATP production (Fig EV1E) and blocked the sperm pseudopodia extension and MO fusion that are induced by SAS (sperm activated subtracts) (Figs EV1C and EV1D). These results suggest that ATP is essential for sperm maturation.

**Fig 1.**
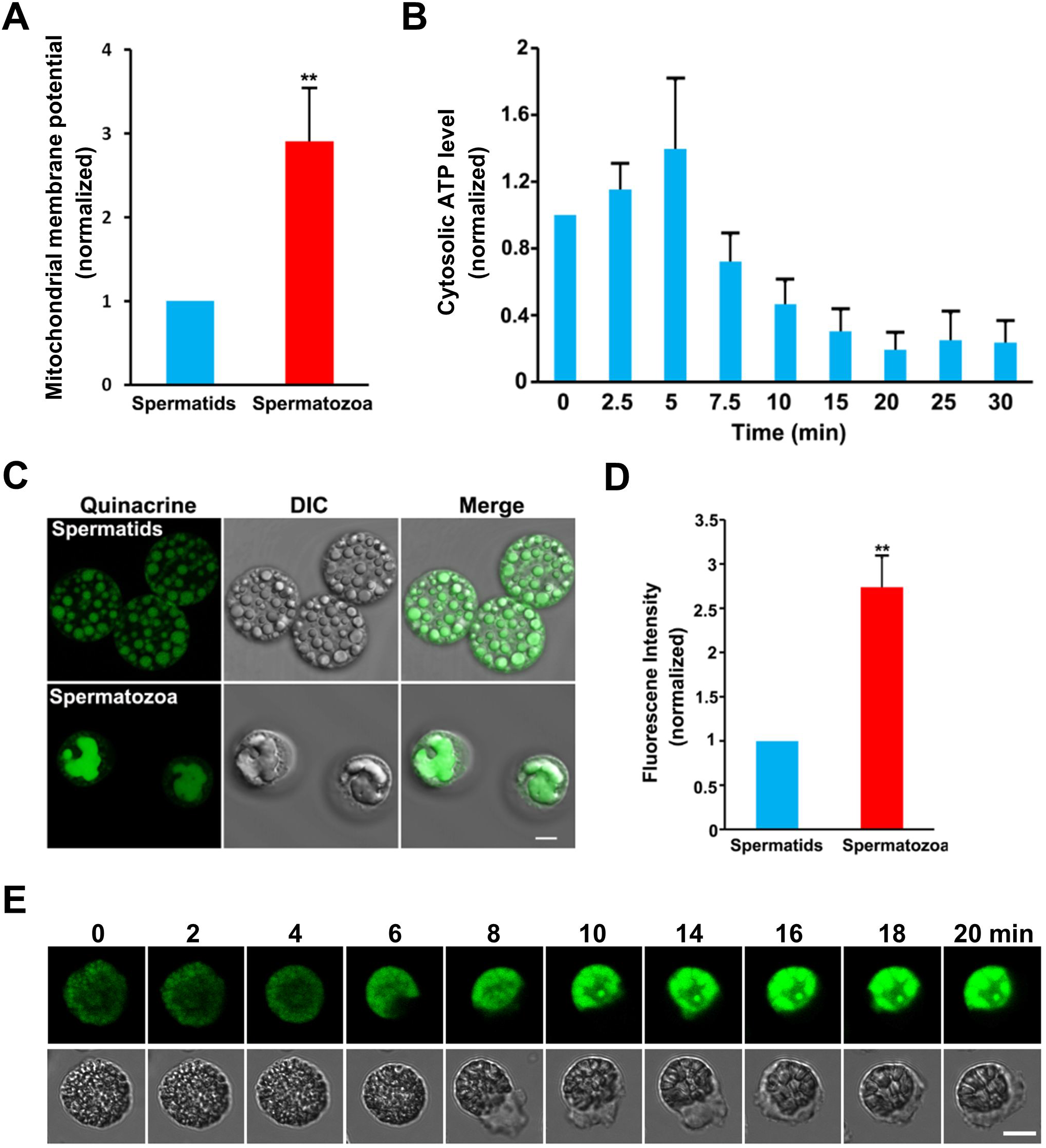
Plentiful ATP is produced during sperm maturation. (A) Statistical results of mitochondrial membrane potential (MMP) in spermatids and spermatozoa were summarized and then normalized. (B) The cytosolic ATP levels were measured with luciferin-luciferase method at a few time points during sperm activation and were normalized with cytosolic ATP concentration at the beginning of measurements. (C) Spermatids and spermatozoa were loaded with quinacrine. ATP was stored in refringent granules (RG) in spermatids and spermatozoa. (D) Quantifying fluorescence intensity of quinacrine. Spermatids and spermatozoa were stained with quinacrine then were analyzed with a flow cytometer. (E) The dynamics of ATP storage and RG fusion during sperm maturation. Values represent means ± SEM. **,*p*<0.01. Bar= 5 μm.

**Fig 2.**
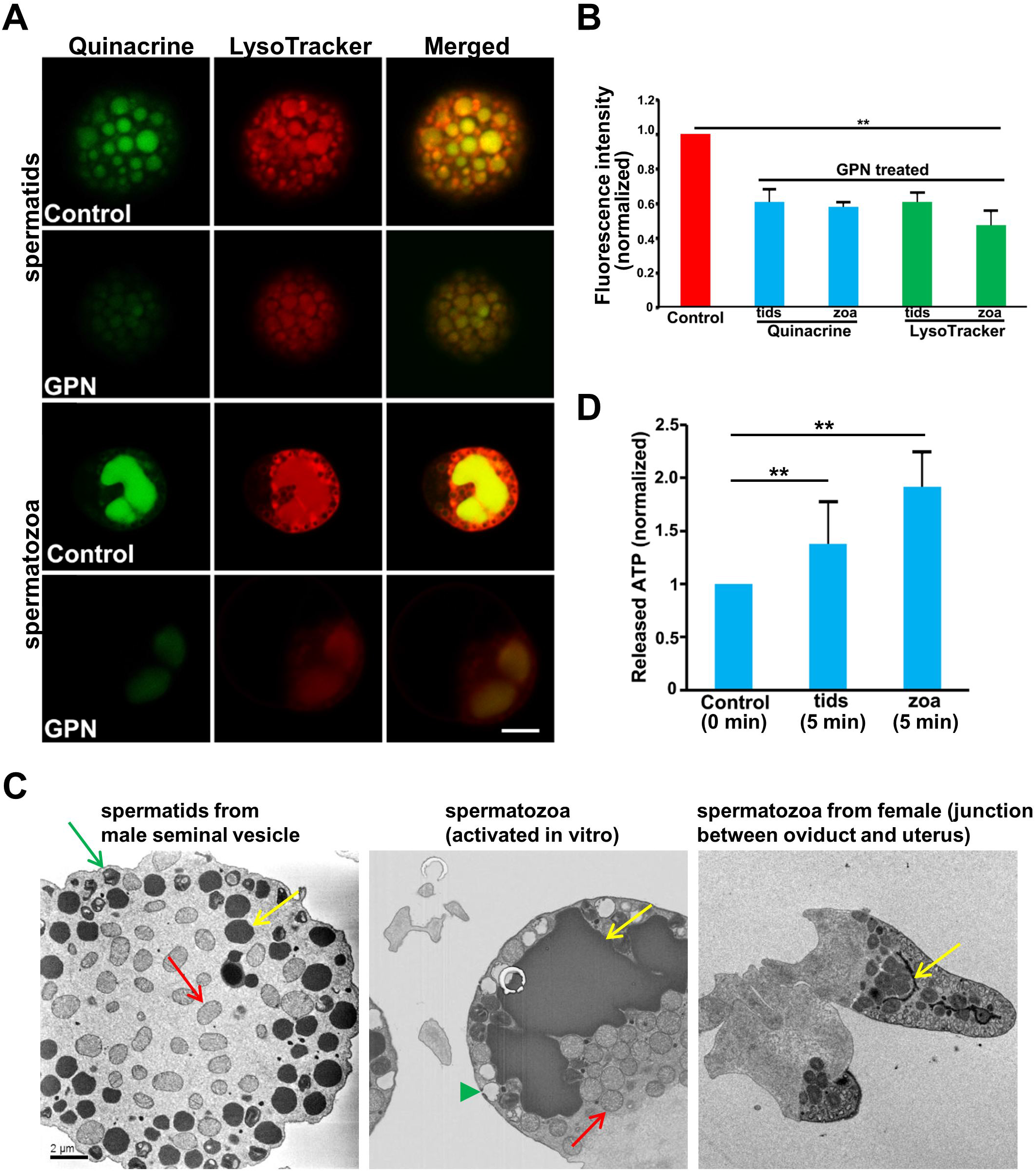
RG is a lysosome-releated and ATP-containing organelle. (A) Both quinacrine and LysoTracker were enriched in no-fused RG in spermatids and fused RG in spermatozoa. Signals of quinacrine and LysoTracker were attenuated by GPN. Bar=5 μm. (B) Fluorescence intensities of quinacrine and LysoTracker in spermatids and spermatozoa were analyzed with flow cytometryand then were quantified. Compared with control, GPN treatment decreased fluorescence intensities of quinacrine and LysoTracker in spermatids and spermatozoa significantly. (C) Transmission electron microscope images show the ultrastructure of spermatid, spermatozoon in vitro and spermatozoa in female reproductive tract. Note that RG show different morphology in these three cell types. RGs are indicated by yellow arrows. Mitochondria are indicated by red arrows. An MO in spermatid is indicated by a green arrow. A fused MO in spermatozoon is indicated by a green arrowhead. (D) Both spermatids and spermatozoa released ATP to extracellular space. In (B) and (D), values represent means ± SEM. **, *p*<0.01.

RGs are specialized organelles that are enriched with amino acids, polypeptides and sugars in *Ascaris suum* sperm (Abbas and Cain, 1981, 1984). During sperm maturation, the small RGs fused with each other and formed a large organelle. Therefore, the sizes and shapes of the RGs were different between spermatids and spermatozoa. Furthermore, the sizes of the RGs in spermatozoa in vitro and in vivo were different. The RGs of spermatozoa in the female reproductive tract disappeared. This indicates that RGs may play a role in sperm fertilization with oocytes (Fig 2C, Movies EV2 and EV3). However, the characteristics and functions of RGs remain unknown. To further characterize ATP storage vesicles in *Ascaris suum* sperm, spermatids and spermatozoa were stained with quinacrine and LysoTracker, a fluorescent dye for labeling acidic organelles. ATP storage vesicles that were visualized with quinacrine could also be labeled with LysoTracker (Fig 2A). This indicated that the RGs were acidic organelles. To further study the characteristics of the RGs, the lysosomotropic compound glycyl-L-phenylalanine 2-naphthylamide (GPN), which is a lysosome-disrupting cathepsin C substrate that causes reversible permeabilization of lysosomes by osmotic swelling, was used to treat sperm. After spermatids and spermatozoa were treated with GPN, the fluorescence intensities of quinacrine and LysoTracker were attenuated significantly (Fig 2A). To study the GPN effect on the storage of ATP and protons, flow cytometry analysis was performed to quantify the fluorescence intensity of sperm treated with or without GPN. The storage of ATP and protons decreased almost 40% in spermatids and spermatozoa upon GPN treatment (Fig 2B). Our data suggest that RG is not only an ATP-containing organelle but also a lysosome-related organelle. Therefore, ATP is stored in acidic organelles in *Ascaris* sperm. This indicates that an appreciable amount of ATP is produced by mitochondria and is stored in RGs.

### Nematode sperm release ATP into the extracellular space via innexin channels

Neural or immune cells are able to release ATP via channels or exocytosis (Guthrie et al., 1999; Junger, 2011; Loiola and Ventura, 2011). Innexins in invertebrates and pannexins in vertebrates mediate ATP release (Dahl et al., 2013; Pinheiro et al., 2013; Samuels et al., 2010; Sandilos et al., 2012). We tested whether nematode sperm release ATP into the extracellular space through innexin channels. First, we measured ATP concentration of sperm medium. Spermatids were washed with buffer, and the ATP concentration in the medium was measured at two time points (0 and the 10th minutes). The ATP concentration in the medium of spermatids (2×10^7^ cells in 500 μL of solution) was 1.93 μM in the beginning and increased to 2.66 μM at the 10th minute. The ATP level in the medium increased, which indicates that nematode spermatids are able to release ATP through autocrine signaling. Second, we tested whether innexins mediated ATP release. Spermatids were treated with carbenoxolone (CBX, 100 μM), which is an innexin channel blocker. CBX attenuated ATP release from 2.66 μM to 1.51 μM at the 10th minute, thereby suggesting that ATP release of spermatids was probably mediated by innexins (Fig 3D). There are 25 members of the innexin family in *Caenorhabditis elegans*, 3 of which are expressed in sperm (Altun et al., 2009). In *Ascaris suum*, transcripts of *innexin-3* and *innexin-11* were detected in the testis (Fig 3A). We raised Innexin-3 antibody to detect its localization on sperm. Innexin-3 (GenBank: ERG82140.1) is composed of 429 amino acids, and its molecular weight is ~50 kDa. In an immunoblotting assay, a single ~50 kDa band was detected in the plasma membrane but not in the cytosol of *Ascaris* sperm. Moreover, these signals in the plasma membrane decreased dramatically when the antibody was preincubated with peptide antigen. No bands were detected when samples were probed with preimmune serum (Fig 3B). In the immunostaining assay, as shown in Fig 3C, Innexin-3 was localized on the plasma membrane of spermatids and spermatozoa. The cytosolic signals were probably nonspecific because they were not blocked by peptide antigen. To further elucidate the role of ATP release in sperm maturation, CBX (100 μM), which was demonstrated to inhibit ATP release, was used, and we found that CBX blocked more than 50% of sperm activation (Fig EV2). This indicates that ATP release that is mediated by innexins plays a role in modulating sperm activation.

**Fig 3.**
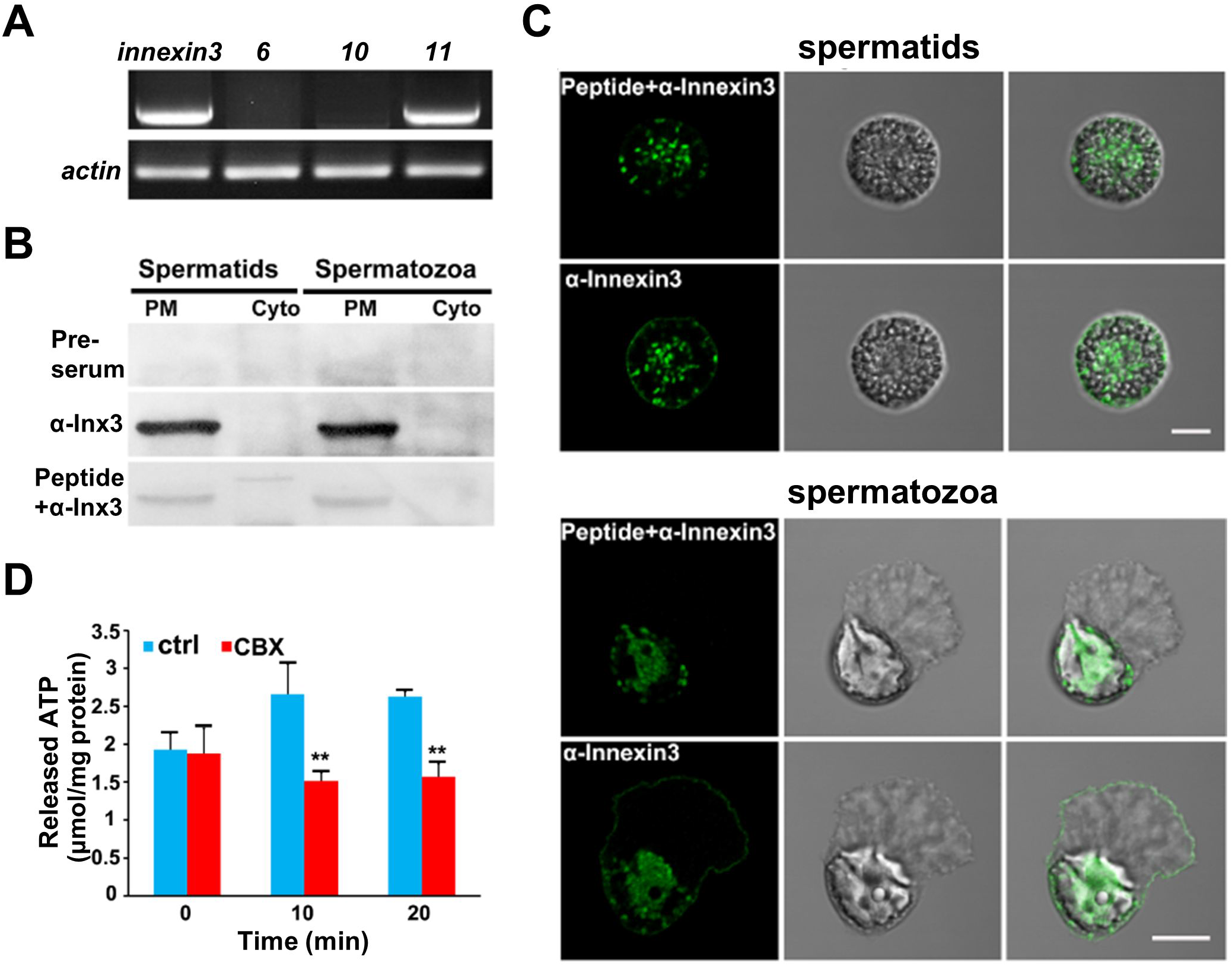
ATP is released through innexin channels. (A) RT-PCR results demonstrated that transcripts of *innexin-3* and *innexin-11* were detected in testis. (B) In Western blotting analysis, Innexin-3 was detected in the plasma membrane fractions of spermatids and spermatozoa with Innexin3 antibody. Peptide antigen and preimmune serum were used to distinguish signals of Innexin-3 from nonspecific signals. Plasma membrane (PM), cytosol (Cyto). (C) Immunofluorescence analysis of Innexin-3 expression in spermatids and spermatozoa. Innexin-3 was expressed in sperm plasma membrane. Peptide antigen was used to distinguish signals of Innexin-3 from nonspecific signals. Bar= 5 μm. (D) ATP levels in medium of spermatids were measured and ATP concentration increased after 10 minutes. CBX, an ATP channel blocker, blocked this process. Values represent means ± SEM. **,*p*<0.01.

### MIG-23, which is a homolog of E-NTPDase, contributes to the ecto-ATPase activity of spermatozoa

To further study the correlation between extracellular ATP and sperm function, we measured the extracellular ATP level at a few time points when spermatids were activated. As shown in Fig 4A, the extracellular ATP level increased by almost 1-fold compared with the control at the 5th minute after SAS was added. This was consistent with the intracellular ATP level (Fig 1B). However, after 5 minutes, the extracellular ATP level decreased gradually. This raises the question of why extracellular ATP is attenuated significantly during sperm maturation but mitochondrial activity is higher in the same time. This implies that released ATP is hydrolyzed by ecto-ATPase on spermatozoa. We hypothesize that ecto-ATPases on the sperm surface are activated during sperm maturation and degrade released ATP. To test this hypothesis, exogenous ATP (5 μM) was added to spermatozoa (10^7^ cells in 500 μl solution). The spermatozoa hydrolyzed 80% of the exogenous ATP (5 μM) within 5 minutes. However, when exogenous ATP (5 μM) was added to spermatids (10^7^ cells in 500 μL solution), they did not hydrolyze the exogenous ATP (Fig 4B). This suggests that ecto-ATPase activity of spermatozoa is gained during sperm maturation. Ecto-nucleoside triphosphate diphosphohydrolase (E-NTPDases), ecto-phosphodiesterases/pyrophosphatases, ecto-5’-nucleotidase and alkaline phosphatases are able to hydrolyze extracellular ATP (Sansom et al., 2008). It has been reported that ZnCl_2_, 4,4’-diisothiocyanatostilbene-2,2’-disulfonic acid (DIDS), reactive blue-2 (RB-2) and suramin decrease mammalian E-NTPDase activity (Baqi et al., 2009; Barros et al., 2000; Dowd et al., 1999; Iqbal et al., 2005; Leal et al., 2005). We tested whether these inhibitors also functioned in *Ascaris suum* sperm. Spermatozoa (2×10^7^ cells in 500 μL solution) were treated with ZnCl_2_ (500 μM), DIDS (10 μM), RB-2 (50 μM) and suramin (10 μM) for 10 minutes. Then, exogenous ATP (5 μM) was added. As shown in Fig EV3, these inhibitors, especially RB-2, significantly impeded exogenous ATP degradation by spermatozoa. These data demonstrate that E-NTPDases contribute to the ecto-ATPase activity of spermatozoa in *Ascaris suum*.

**Fig 4.**
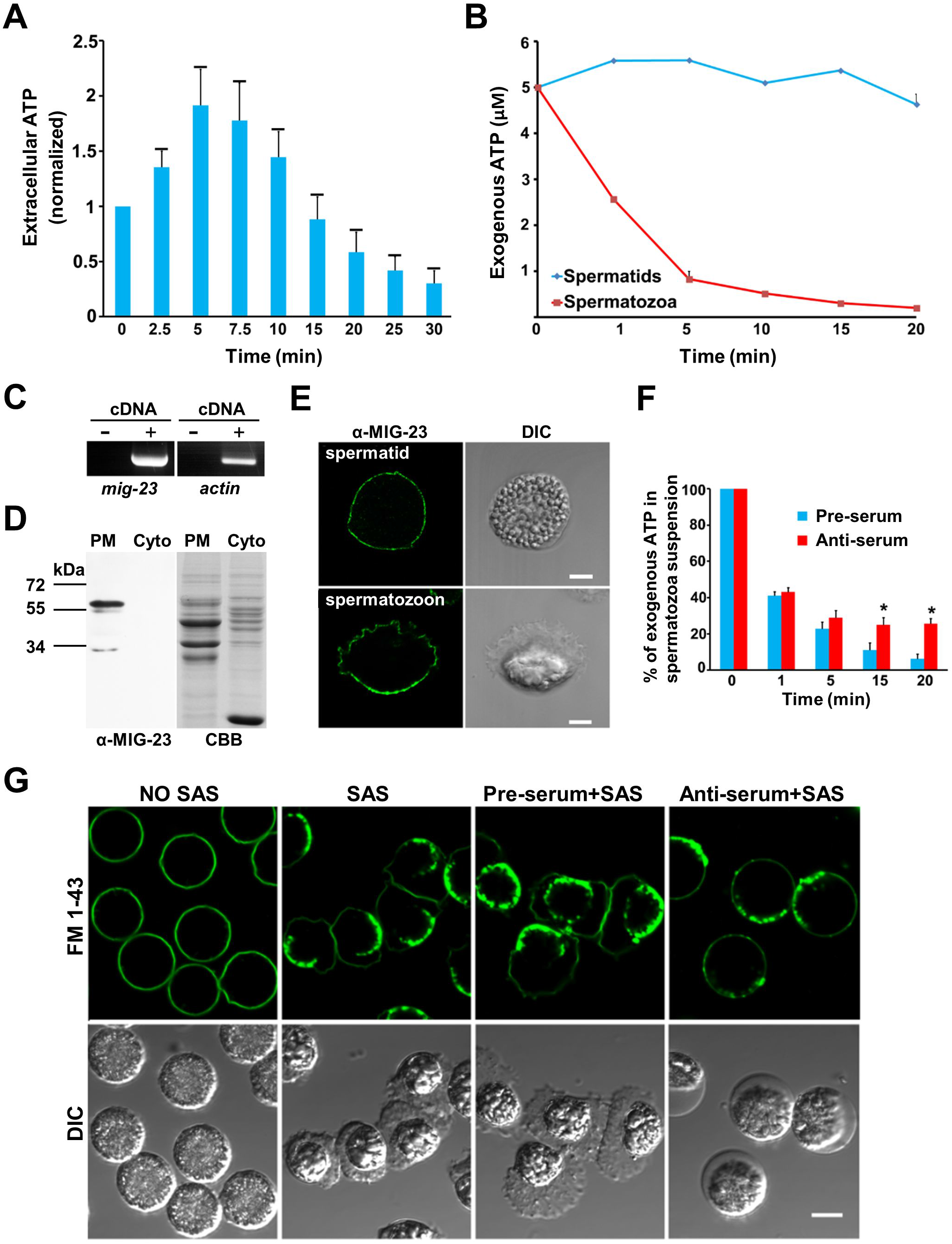
MIG-23 is required for sperm function. (A) Extracellular ATP levels at a few time points were measured after spermatids were activated. Extracellular ATP level increased in 5 minutes and then decreased gradually. (B) Spermatozoa showed ecto-ATPase activity and degraded exogenous ATP. However, spermatids did not degrade exogenous ATP. (C) Transcript of *mig-23* was detected in testis. (D) Anti-MIG-23 antibody detected the signal in the plasma membrane (PM) but not in cytosol (cyto) in immunoblotting assay (left panel). PM and cytosol fractions were analyzed by SDS-PAGE (right panel). (E) In immunostainning assay, non-permeabilized spermatids and spermatozoa were labeled by anti-MIG-23 antibody. MIG-23 was localized on the plasma membrane. (F) MIG-23 antiserum partially attenuated the process in which exogenous ATP (5 μM) was degraded by spermatozoa. (G) Sperm treated with MIG-23 antiserum showed normal MO fusion indicated by punctate signals of FM 1-43, a membrane probe, during sperm maturation. However, these sperm formed round pseudopodia in which MSP-based filaments were not maintained. As a control, sperm treated with preimmune serum showed normal MO fusion and pseudopodia. In (A), (B), and (F), values represent means ± SEM. *, *p*<0.05. In (E) and (G), bar=5 μm.

Mammalian E-NTPDases 1, 2, 3 and 8 localize on the plasma membrane. They regulate multiple and diverse physiological processes in cells, such as neutrophil chemotaxis, T-cell activation and set points of cardiac fibroblasts (Corriden et al., 2008b; Knowles, 2011; Lu and Insel, 2013; Robson et al., 2006). We performed BLAST with the human E-NTPDase 2 sequence and obtained the homolog nucleoside-diphosphatase MIG-23 (GenBank: ERG84046.1) in *Ascaris suum*. MIG-23 has 551 amino acids, and its molecular weight is ~60 kDa. It shares 41% similarity with mammalian E-NTPD1, 42% with E-NTPD2, 42% with E-NTPD3, and 41% with E-NTPD8. MIG-23 is conserved in various species. The alignments of MIG-23 with E-NTPD2 in *C. elegans, Homo sapiens, Mus musculus*, and *Xenopus tropicalis* are shown in Fig EV4. The transcription of MIG-23 was detected in the testis (Fig 4C). MIG-23 peptide antibody was raised to investigate the precise localization of MIG-23 in sperm. In an immunoblotting assay, a band of ~60 kDa was detected in the plasma membrane extracts but absent in the cytosolic extracts (Fig 4D). In immunofluorescence assays, when sperm were not permeabilized, MIG-23 signals were detected on the plasma membrane (Fig 4E). The N-terminus and C-terminus of MIG-23 are in the cytosol. MIG-23 has two transmembrane domains and a large extracellular loop. To test whether the functional domain of MIG-23 is present on the surface of sperm, MIG-23 antiserum (1:100 dilution) was incubated with spermatozoa to block MIG-23 activity. The MIG-23 antiserum significantly attenuated the ability of spermatozoa to hydrolyze exogenous ATP (5 μM) at the 10th and the 15th minutes (Fig 4F). This indicates that the functional domain of MIG-23 localizes to the outer plasma membrane of spermatozoa and that its activity can be blocked by MIG-23 antiserum.

### MIG-23 is required for MSP-based filament assembly and sperm migration

To further evaluate the role of MIG-23 in sperm function, MIG-23 activity was blocked by MIG-23 antiserum (1:100 dilution). Spermatids that were preincubated with MIG-23 antiserum were not activated by SAS normally although MOs fused with the plasma membrane. These sperm showed round pseudopodia without leading edge protrusions when they were activated in a few minutes (Fig 4G), thereby indicating abnormal MSP filament assembly and disassembly in these sperm. To examine the effects of MIG-23 on MSP filament dynamics, we captured images of MSP filament dynamics in vitro. Blocking MIG-23 activity led to increased MSP filament disassembly in pseudopodia. This caused the MSP filament cytoskeleton to fail to be maintained and the MSP filaments to be depolymerized in a few minutes. The balance between disassembly and assembly of MSP filaments was destroyed because of the lack of MIG-23 activity. Nematode spermatozoa migration is based on the MSP filament cytoskeleton. Cytoskeleton assembly and disassembly are tightly coupled with sperm migration (Bottino et al., 2002; Wolgemuth et al., 2005). Thus, MIG-23 activity is required for promoting MSP filament assembly. Spermatozoa that were pretreated with MIG-23 antiserum were not able to migrate, although vesicular exocytosis occurred. Blocking MIG-23 activity with MIG-23 antiserum caused MSP-based filaments to be maintained in a short time. This leads to spermatozoa not to migrate(Fig 4G, Movies EV4 and EV5). In cell-free system, fiber pretreated with Mig-23 antiserum grew slower compared with control ( Movies EV6 and EV7). Together, these data indicate that the membrane protein MIG-23 plays an essential role in regulating sperm migration. Cytoskeletal dynamics are modulated precisely by factors such as MSP-associated proteins, protein phosphorylation and dephosphorylation (Yi et al., 2009). MIG-23, which is a membrane protein of which the functional domain localizes on the outer membrane, regulates sperm MSP filament dynamics and migration by modulating extracellular ATP levels. Purinergic Signaling mediated by Mig-23 is important for *Ascaris* sperm maturation.

## Discussion

Previous work demonstrated that the MSP-based cytoskeleton is assembled, pseudopodia form, membranous organelles fuse with the plasma membrane and granular organelles fuse with each other during sperm maturation in *Ascaris suum*. In addition, we found that MMP increased significantly and that a large amount of ATP was produced to satisfy physiological needs during sperm maturation. The produced ATP was rearranged by sperm efficiently. Otherwise, abundant ATP in the cytosol might have been unfavorable for sperm maturation. Some ATP was stored in RGs, although the mechanism of ATP accumulation in RGs remains unclear. To our knowledge, this is the first study demonstrating that RGs are ATP-containing vesicles. Some ATP was released into the extracellular space through innexin channels that were localized on the plasma membrane of sperm. Therefore, nematode sperm are able to release ATP into the extracellular space. ATP release in astrocytes modulates depressive-like behavior in mice. ATP release is mediated by calcium and V-ATPase (Cao et al., 2013; Coco et al., 2003). Exogenous ATP treatment of human sperm improved in vitro fertilization (IVF) results by increasing sperm motility (Rodríguez-Miranda et al., 2008). In the nematode *Ascaris suum*, we identified that ATP release played a role in sperm activation. In some species, ATP is released through ATP channels and exocytosis. Pannexin in vertebrates and innexin in invertebrates mediate ATP release (Lu et al., 2015). Pannexin-3, which is one of three members of the pannexin family, is present in the epididymis of adult rats. It might play a role in the maturation and transport of sperm (Turmel et al., 2011). The function of INX-14 in the nematode *C. elegans* is essential for guiding spermatozoa to oocytes (Edmonds et al., 2011). We found that nematode *Ascaris suum* sperm were able to release ATP through Innexin-3 and Innexin-11. The mRNAs of *innexin-3* and *innexin-11* were expressed in *Ascaris suum* testis, and the Innexin-3 protein was localized on the plasma membrane of sperm. Carbenoxolone (CBX), which is a blocker of ATP channels, inhibited ATP release and attenuated sperm maturation. This indicates that ATP is released via ATP channels and that ATP release is required for sperm function.

Released ATP is not stable and is hydrolyzed by E-NTPDases (Lu and Insel, 2013). This process facilitates various cell functions. For example, extracellular ATP hydrolysis blocks synaptic transmission (Vroman et al., 2014). *Ascaris* spermatozoa show ecto-ATPase activity. MIG-23 in *Ascaris suum*, which is a homolog of human E-NTPDases that is localized on the sperm surface, contributes to the ecto-ATPase activity of spermatozoa. Human membrane E-NTPDase contains an active site that is exposed to the extracellular space (Chiang and Knowles, 2008). Similar to human membrane E-NTPDase, the functional domain of MIG-23 is also exposed to the extracellular space. The ecto-ATPase activity of spermatozoa decreased partially when MIG-23 activity was blocked by MIG-23 antibody. Spermatozoa that were pretreated with MIG-23 antiserum showed migration defects because MSP filament assembly was slower than disassembly. This led to depolymerization of the MSP filament cytoskeleton. Taken together, we conclude that MIG-23 is involved in sperm function by modulating extracellular ATP levels. MIG-23 is important for sperm migration and is a newly identified component of fertility pathways in *Ascaris suum*.

## Materials and Methods

### Worm harvest and sperm preparation

*Ascaris suu*m male worms were collected from slaughterhouse and kept in worm buffer (PBS containing 100 mM NaHCO_3_, pH 7.0, 37 °C). Sperm (spermatids) were harvested from seminal vesicles together with seminal fluid into HKB buffer (50 mM HEPES, 70 mM KCl, 10 mM NaHCO_3_, pH 7.6 for stock, adjust pH to 7.1 with CO_2_).

### Measurement of mitochondrial membrane potential (MMP)

MMP of sperm was measured with JC-1 kit (Beyotime). Sperm were stained with JC-1 (5 μg/ml) for 20 min at 37 °C. According to the instruction, after being washed twice with washing buffer, sperm were analyzed with flow cytometry. The parameters were set as λex 488 nm and λem 530 nm for JC-1 monomer, λex 525 nm and λem 590 nm for JC-1 aggregates.

### Measurement of ATP concentrations

ATP concentration was measured using ATP assay kit (Beyotime). Luciferin oxidation was catalyzed by firefly luciferase in the presence of ATP. Luminescence emitted was detected with a Sirius luminometer (Berthold Detection Systems). In measurements of extracellular ATP, spermatids were washed twice. At the indicated times, sperm suspensions (50 μL) were collected and centrifuged at 12,000 rpm for 1 min to separate sperm from medium. The concentrations of extracellular ATP in medium could be measured immediately. To detect cytosolic ATP concentrtion, sperm were lysed with lysis buffer in the kit and centrifuged at 13,000 rpm for 10 min at 4 °C to remove debris. The supernatants were collected for cytosolic ATP measurement. The protein concentrations in the sperm lysate was determined using BCA protein assay kit (Beyotime) and were utilized for normalization.

In spermatozoa degrading exogenous ATP assay, 5 μM ATP was added into spermatozoa suspensions (HKB buffer as control) and at the indicated time 50 μL samples were collected and centrifuged at 12,000 rpm for 1 min. Concentrations of ATP in supernatants were measured to indicate amounts of remaining exogenous ATP. To test the effects of ecto-ATPase inhibitors, spermatozoa were pr-incubated with inhibitors of ecto-ATPase (ZnCl_2_ 250 μM, DIDS 5 μM, RB-2 50 μM, suramin 10 μM) for 10 min (20 min for MIG-23 antiserum and preimmune serum) at 37 °C, then 5 μM exogenous ATP was added.

### Blockage of sperm activation triggered by SAS

In HKB buffer spermatids were pre-incubated with reagents (CBX 100 μM, CCCP 10 μM, antimycin A 10 μM, oligomycin 10 μM) for 10 min (20 min for MIG-23 antiserum and preimmune serum diluted at 1:100) at 37 °C then SAS was added to activate sperm. Sperm were stained with FM 1-43 (5 μg/ml) for 5 min and then observed under an FV1200 confocal microscope equipped with a 60×/1.35 NA oil immersion objective (Olympus) (λex 488 nm and λem 505 nm) to detect MO fusion.

### Sub-cellular fraction of sperm

Sub-cellular fraction was performed according to protocol from Dr. Ren (Ren et al., 2006). Sperm harvested were washed twice with HKB buffer then were resuspended in 5 mL lysis buffer (20 mM HEPES, 10 mM KCl, 1.5 mM MgCl_2_, 1 mM EDTA, 1 mM EGTA, 1 mM DTT, 0.1 mM PMSF, 1×protease inhibitor cocktail, 250 mM sucrose) and incubate on ice for 30 min. After 30 strokes of Dounce homogenization, the homogenate was layered onto 5 mL lysis buffer (1 M sucrose) and centrifuged at 6,000 g for 10 min at 4 °C. Upper phase (cytoplasmic fraction) was transferred and centrifuged twice at 9,800 g for 10 min at 4 °C. Then the supernatant was collected and centrifuged at 100,000 g for 30 min at 4 °C. The supernatant obtained was S-100 media (cytosol without mitochondria) and the pellet was plasma membrane.

### RT-PCR assay

cDNA was synthesized from total RNA in testis of *Ascaris suum*. Primers in PCR: *innexin-3*, F, 5’-CTCGAGATGTTCCTCGGTATCCCACAACTGA-3’ and R, 5’-G GTACCACGCGGTAACTCTTTCATCGGCAGTG-3’; *innexin-6*, F, 5’-AAGCTTATGAGTTC ACAAATTGGCGCAATCG-3’ and R, 5’-GGATCCGACGGCTTTATTTCCTTTCTGGAGC-3’; *innexin-10*, F, 5’-AAGCTTATGGTGCTCACAACGGTCCTTTCAA-3’ and R, 5’-GGATCCG ATGATAAGATTTGGGACCTTCTTTGGCG-3’; *innexin-11*, F, 5’-CTCGAGATGATGATCGAAAGTCTCATGGCGA-3’ and R, 5’-GGTACCGTCATCGGAAGGTGTTTTCGGTGAG-3’; *mi g-23*, F, 5’-GCGGCCGCATGGTGCGAGGCATGCTGAG-3’ and R, 5’-GGATCCGAAAAGC TTGGTATATTGTAAT-3’; *actin*, F, 5’-AATCAAAGCGAGGTATCCTCAC-3’ and R, 5’-TGG GTCATCTTTTCTCTGTTTG-3’.

### Immunostaining assay

Both Innexin-3 polypeptide antibody and MIG-23 polypeptide antibody were raised in rabbit. Peptide sequence of Innexin-3 was PDEISRPLSALQGDPIDD. Peptide sequence of MIG-23 was RDVQRRYLLDKRR. Both antibodies were purified. Western blotting was performed according to standard method. Both Innexin-3 antibody and MIG-23 antibody were diluted at 1:500. In immunofluorescence assay, sperm were fixed with 1.25% glutaraldehyde in HKB buffer and then were permeabilized with 0.5% Triton X-100 or not permeabilized. Sperm were incubated with blocking solution (2% BSA in PBS) for 4 h at room temperature and then were stained with primary antibody (anti-Innexin-3, 1:50; anti-MIG-23, 1:50) overnight at 4°C. After washing with PBS, secondary antibodies (anti-rabbit IgG–Alexa Fluor 488, 1:400) were incubated with sperm for 2 h at room temperature. Samples were observed under a confocal microscope (63×/NA 1.4 oil objective, Carl Zeiss) with 488/543 nm laser lines.

### Transmission Electron microscopy

All the sperm were fixed with GTS-Fixative (2.5% glutaraldehyde, 2 mg/mL tannic acid and 0.5 mg/mL saponin in HKB) for 40 min on a thermanox plastic coverslip (EMS, USA), followed by washing in HKB buffer and then water. They were post-fixed in 1% osmium tetroxide for 30 min, dehydrated in a graded series of ethanol followed by propylene oxide, and then infiltrated and embedded with EMbed-812 resin (EMS, USA). Ultrathin sections (80 nm) were cut on a Leica UC6 ultramicrotome, collected on formvar-coated copper grids and stained with uranyl acetate and lead citrate. TEM images were captured using an FEI Spirit 120 kV electron microscope (FEI, USA) operated at 100 kV.

### Preparation of sperm extracts for MSP fiber assay *in vitro*

Spermatids were treated with preimmune serum or MIG-23 antiserum (1:100 dilution) for 20 min at 37 °C and then SAS together with preimmune serum or MIG-23 antiserum was added to activate sperm. Sperm were harvested after 10 min and were centrifuged at 12,000 rpm for 1 min. After removing the supernatant, the cell pellets were frozen at −80°C overnight and then thawed on ice. Sperm were centrifuged at 23,000 rpm for 15 min at 4°C and then the supernatant was collected and centrifuged at 100,000 g for 1 hr at 4°C. The supernatant (S100) was used for the fiber assay. In this assay, S100 was diluted at 1:5 in KPM buffer (0.5 mM MgCl_2_, 10 mM potassium phosphate, pH 6.8) and 1 mM ATP was added. After 10 min, the solutions were pipetted into glass chambers and examined on an Axio Imager M2 microscope (Carl Zeiss) equipped with a 40×/NA 0.95 Ph3 Korr objective. Images were acquired using a Zeiss Axiocam digital camera and were analyzed using the MetaMorph software.

## Acknowledgements

This work was supported by grants from the National Natural Science Foundation of China (32070694, 31872822, 31571436 to Y.Z.) and the National Key Research and Development Program of China (No. 2018YFC1003500).

## Abbreviations used

E-NTPDase: Ecto-nucleoside triphosphate diphosphohydrolase
MSP: Major sperm protein
MO: Membranous organelle
RG: Refringent granule
ATP: Adenosine-5'-triphosphate

## Author contributions

Qiushi Wang and Ruijun He: investigation, data curation, and writing original draft. Qi Zhang and Jin Shan: methodology, formal analysis, and validation. Yanmei Zhao: editing and project administration. Xia Wang: conceptualization, writing, reviewing, and supervision.

## Figure Legends

**Fig EV1.**
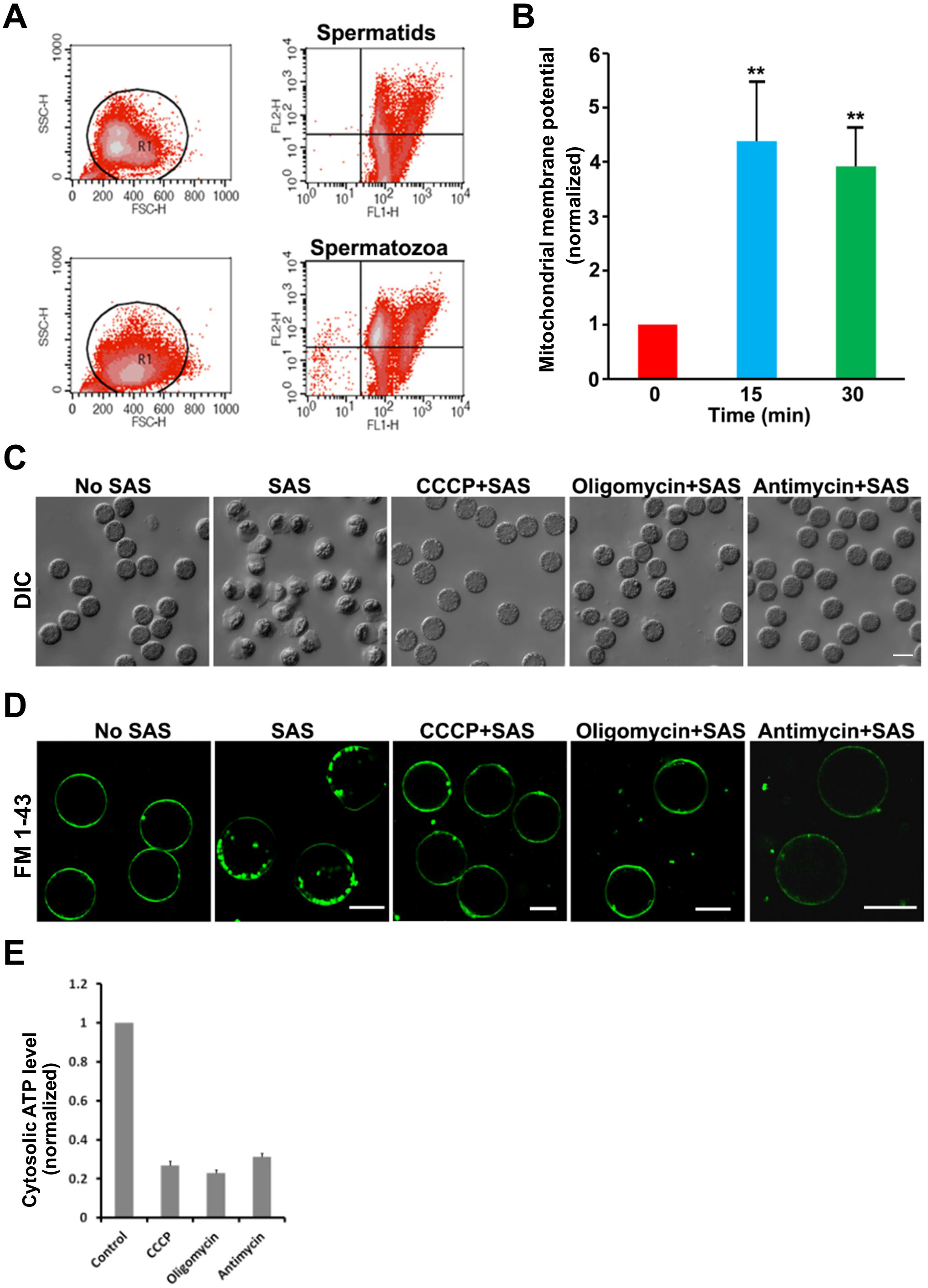
Mitochondrial activity is essential for sperm activation. (A) Spermatids and spermatozoa loaded with JC-1, an indicator of mitochondrial membrane potential (MMP), were analyzed by flow cytometry. (B) MMPs of spermatids, sperm activated at 15 and 30 min were measured and normalized. (C) Sperm treated by CCCP (10 μM), antimycin A (10 μM), and oligomycin (10 μM) were not activated by SAS. These reagents blocked pseudopodia extension in sperm. (D) CCCP, oligomycin, and antimycin A inhibited MO fusion with the plasma membrane indicated by FM 1-43, a membrane probe, during sperm maturation. In (C) and (D), bar=5 μm. (E) Cytosolic ATP levels in sperm were measured when they were treated by CCCP, antimycinA and oligomycin. Values represent means ± SEM. **, *p*<0.01.

**Fig EV2.**
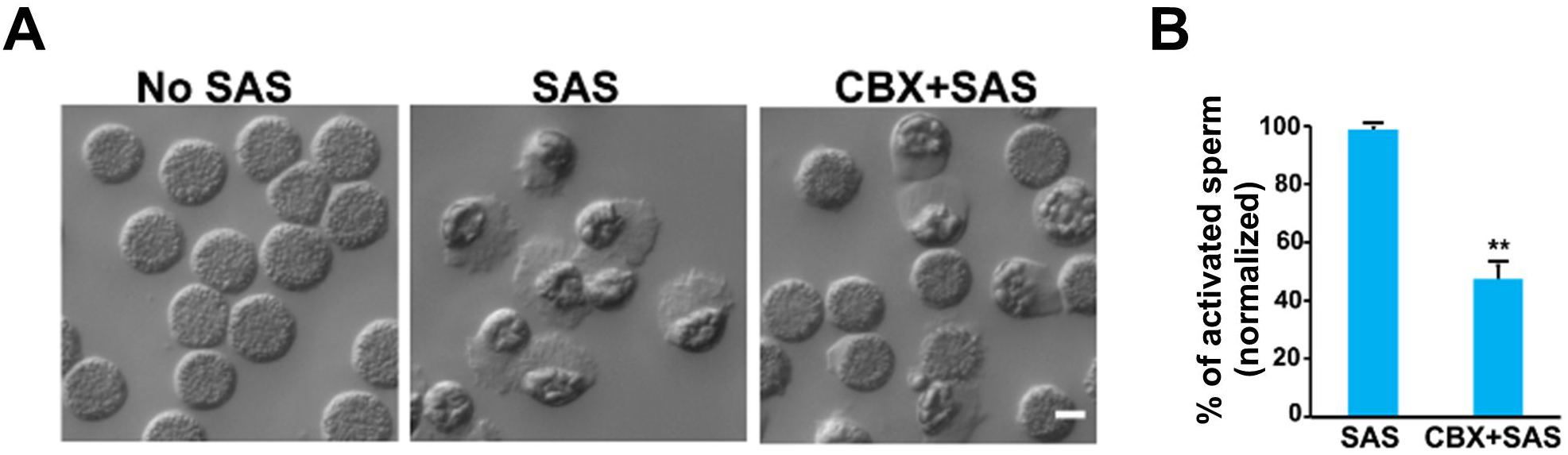
ATP release is required for sperm activation. (A) Some spermatids were not activated normally by activator (SAS) when they were treated by CBX. (B) Quantifying effects of CBX on sperm activation. CBX blocked around 50% of sperm activation (no pseudopodia extension and MSP filaments assembly). Values represent means ± SEM. **, *p*<0.01. Bar=5 μm.

**Fig EV3.**
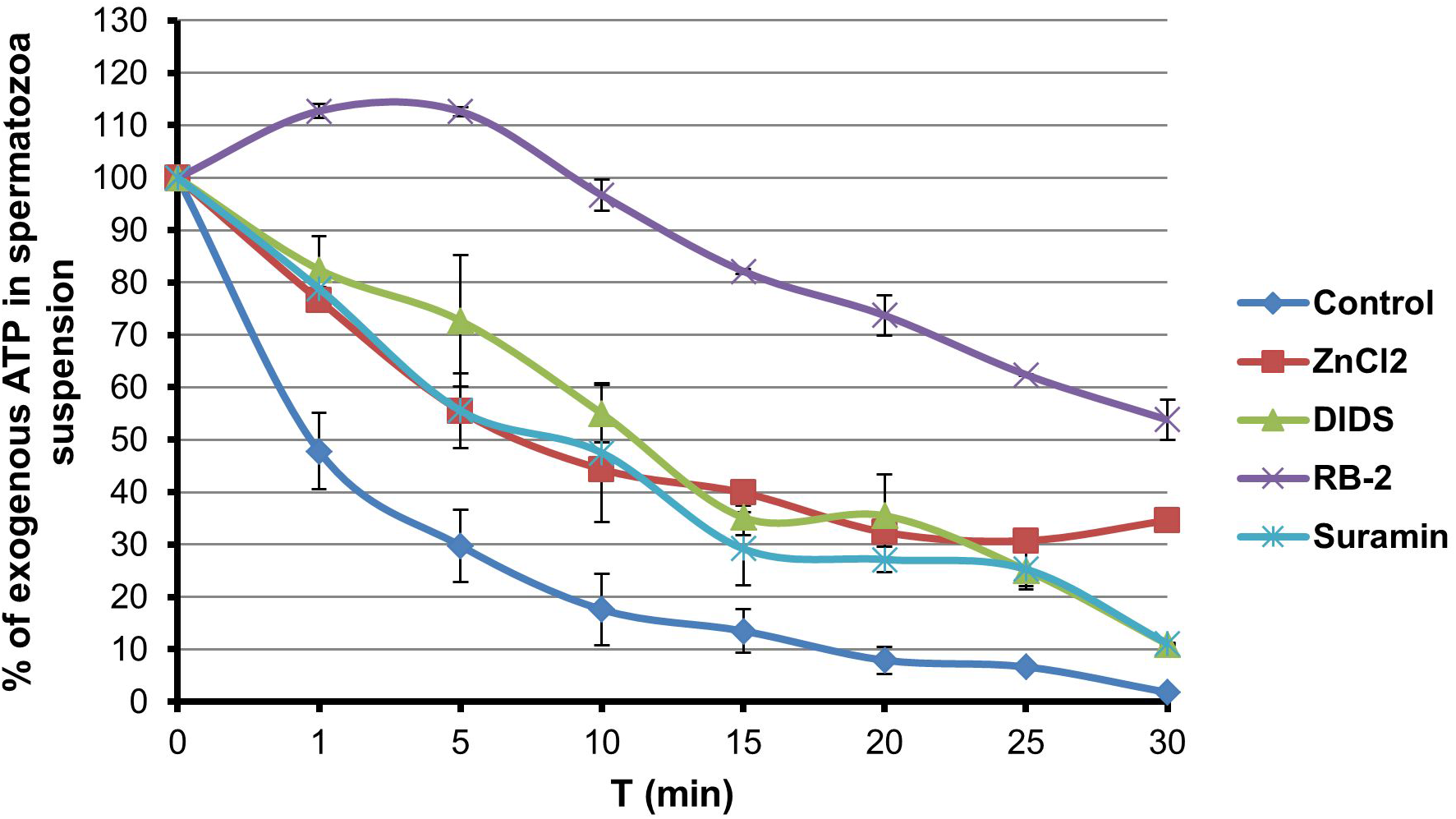
Spermatozoa shows ecto-ATPase activity. Exogenous ATP degradation mediated by spermatozoa was attenuated by inhibitors of ecto-ATPases (ZnCl_2_ 250 μM, DIDS 5 μM, RB-2 50 μM, suramin 10 μM). 5 μM exogenous ATP (set as 100%) was added into HKB buffer with spermatozoa.

**Fig EV4.**
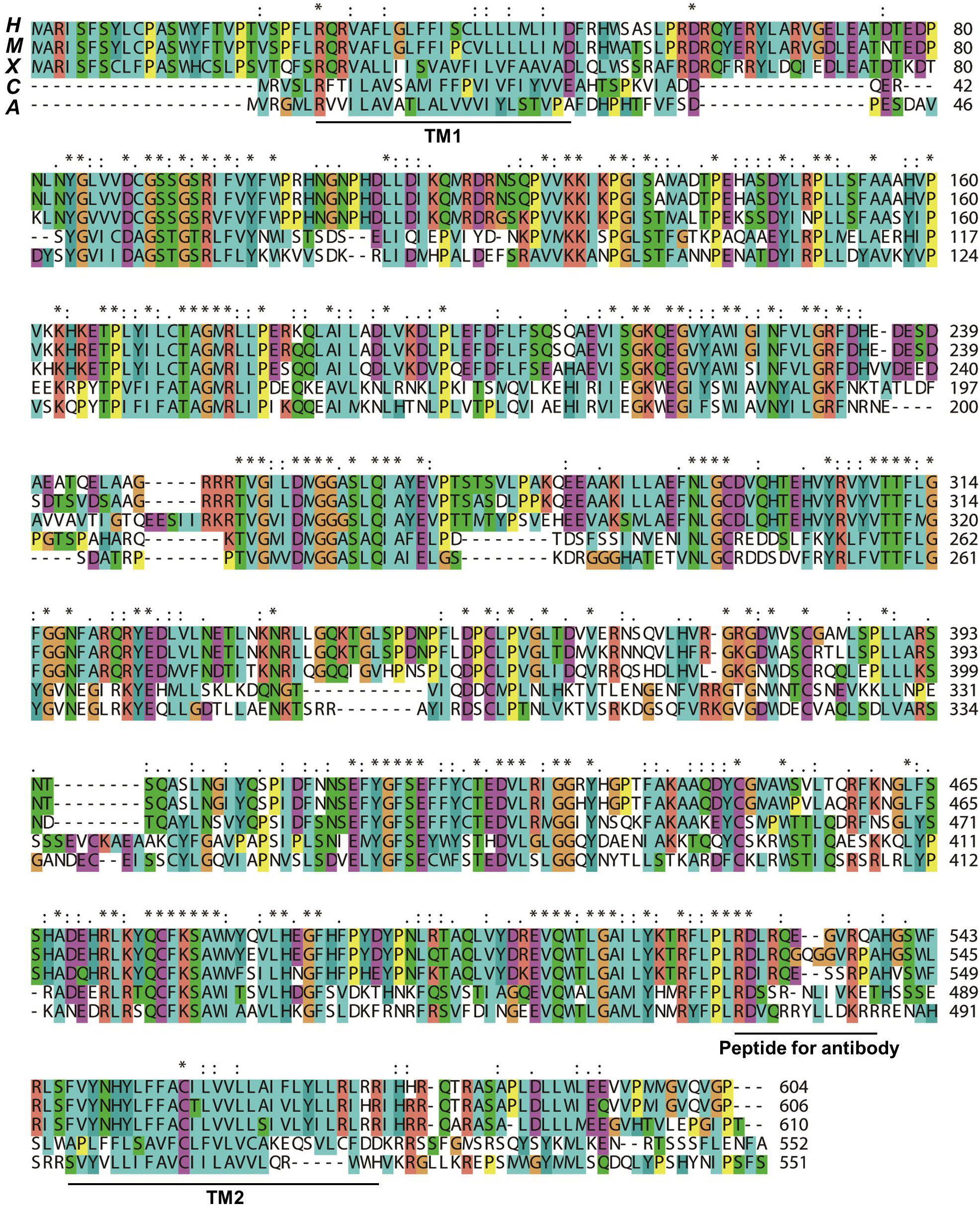
Alignment of MIG-23 in *Ascaris suum*. Alignment of MIG-23 in *Ascaris suum* (A) with E-NTPDases in *C. elegans* (C), *Homo sapiens* (H), *Mus musculus* (M), and *Xenopus tropicalis* (X). TM1 and TM2 (black line) represent transmembrane domains of MIG-23. The amino acids between TM1 and TM2 form a big loop which is functional domain of MIG-23. The amino acid sequence which was used to raise polypeptide antibody is indicated.

**Movie EV1. Dynamic ATP storage during sperm maturation.** Spermatids were loaded with quinacrine (5 μM) at 37 °C for 10 min, and then sperm maturation was recorded under a confocal microscope.

**Movies EV2 and EV3. Ultrastructure of spermatid and spermatozoon.** RGs are darkly stained small granules dispersed in spermtid (Movie EV2). RGs fuse with each other and form a single and huge organelle in spermatozoon (Movie EV3).

**Movies EV4 and EV5. Blocking activity of MIG-23 inhibits the elongation of MSP fibers *in vitro*.** MSP fiber elongated in the *in vitro* system in which extract of sperm treat with preimmune serum was added (Movie EV4). However, the elongation of MSP fiber was slower when extract of sperm treated with MIG-23 antiserum was added (Movie EV5).

**Movies EV6 and Movie EV7. Blocking activity of MIG-23 inhibits spermatozoa migration.** Spermatozoa treated with preimmune serum (1:100 dilution) showed normal MSP-based cytoskeleton and spermatozoa migrated normally (Movie EV6). Spermatozoa treated with MIG-23 antiserum (1:100 dilution) showed round pseudopod and defective migration (Movie EV7).

